# Membrane activity of a DNA-based ion channel depends on the stability of its double-stranded structure

**DOI:** 10.1101/2021.07.11.451603

**Authors:** Diana Morzy, Himanshu Joshi, Sarah E. Sandler, Aleksei Aksimentiev, Ulrich F. Keyser

**Affiliations:** Cavendish Laboratory, University of Cambridge, JJ Thomson Avenue, Cambridge, CB3 0HE, United Kingdom; Department of Physics, University of Illinois at Urbana–Champaign, 1110 West Green Street, Urbana, Illinois 61801, United States; Beckman Institute for Advanced Science and Technology, University of Illinois at Urbana-Champaign, 405 North Mathews Avenue, Urbana, Illinois 61801, United States

**Keywords:** DNA structures, lipids, membranes, tilt, nicks, pore-forming, protein-mimicking, synthetic ion channel

## Abstract

Structural DNA nanotechnology has emerged as a promising method for designing spontaneously-inserting and fully-controllable synthetic ion channels. However, both insertion efficiency and stability of existing DNA-based ion channels leave much room for improvement. Here, we demonstrate an approach to overcoming the unfavorable DNA-lipid interactions that hinder the formation of a stable transmembrane pore. Our all-atom MD simulations and experiments show that the insertion-driving cholesterol modifications, when introduced at an end of a DNA strand, are likely to cause fraying of the terminal base pairs as the DNA nanostructure adopts its energy-minimum configuration in the membrane. We also find that fraying of base pairs distorts nicked DNA constructs when embedded in a lipid bilayer. Here, we show that DNA nanostructures that do not have discontinuities (nicks) in their DNA backbones form considerably more stable DNA-induced conductive pores and insert into lipid membranes with a higher efficiency than the equivalent nicked constructs. Moreover, lack of nicks allows to design and maintain membrane-spanning helices in a tilted orientation within lipid bilayer. Thus, reducing the conformational degrees of freedom of the DNA nanostructures enables better control over their function as synthetic ion channels.

## Introduction

Due to the unparalleled ease of controlling nucleic acids’ structure at the molecular scale, DNA nanoengineering is widely investigated for variety of applications, amongst them mimicking membrane remodelling proteins^1,2^, biological sensing^3,4^, building nanopores^5–7^, designing plasmonic architectures^8^. Alongside these implementations, synthetic biology aims to create DNA-based transmembrane structures working as artificial enzymes and ion channels^9–14^. Such constructs can potentially form a vast library of treatment platforms, acting as drug delivery systems or substituting damaged natural proteins^15–18^. However, the negatively charged DNA backbone and the resulting hydrophilicity prevent spontaneous membrane spanning. To circumvent the unfavorable DNA-lipid interactions, membrane protein mimics are most commonly modified with hydrophobic moieties, particularly cholesterol^11,19–23^.

For simple DNA architectures to span membranes, hydrophobic anchors need to be placed on the opposite sides of the structure. Their strong membrane affinity can then force the nucleic acid to cross the bilayer, since no other conformation that would allow cholesterols’ insertion is possible. Cholesterol-modified DNA will *attach* to a membrane, but only when it is pulled with opposing cholesterol molecules will DNA *insert* into the bilayer. Building of biological nucleic acid nanopores is thus based on designing such architectures, with hydrophobicity distributed in a way that forces the hydrophilic DNA to span the membrane^10,19,22,24^. Aiming at finding a comprehensive description of design principles, numerous variables are tested, including shape^24,25^, number and position of hydrophobic modifications^10,13,26^ and cholesterol-mediated clustering^27^. However, a less-studied aspect of the DNA nanostructure design is also playing an important role: its base pairing stability and backbone integrity.

The repulsive DNA-lipid interactions introduce a strain in the helices: if the structure unwinds, cholesterols will not remain on the opposite sides of the double-stranded structure, and in turn DNA will not cross the membrane. The ion channel activity resulting from membrane-spanning is therefore strongly dependent on the stability of the base pairs in the double helix and the resulting positions of all hydrophobic modifications.

Even though the unwinding is prevented by DNA strands’ complementarity, stable formation of all base pairs should not be taken for granted. Particularly, DNA duplexes are subject to fraying in close proximity of nicks - discontinuities in the sugar-phosphate backbone of one of the strands forming the double-helix. Base stacking interactions are less stable and transient resulting in switching between stacked (straight) and unstacked (kinked) conformations^28–30^. The consequence is a reduction of DNA stability near a nick in the backbone.

Importantly, hydrophobic modifications are often introduced at such discontinuities^10,11,13,31^ due to the relative ease of chemical modification. The free energy gain of a cholesterol molecule’s insertion in a lipid membrane is about – 75 kJ/mol^32,33^, while the free energy of a single base pair dissociation ranges from around 6 kJ/mol (AT) to 8 kJ/mol (GC)^34^. The difference between the free energies illustrates why a short stretch of a DNA duplex can be distorted to facilitate the cholesterol insertion, while allowing the hydrophilic DNA backbone to remain outside the hydrophobic bilayer’s core. Broken base pairs at the nick will result in flexing of the nucleic acid domain, in turn distorting the designed structure of the DNA channel. When subject to forces within the bilayer, even carefully designed DNA nanostructure can prove inefficient, due to its relatively unconstrained deformation *in situ*.

Here, we show that placing the hydrophobic modifications at the terminals of a DNA nanostructure indeed affects the functionality of the DNA-membrane assembly. Employing a minimalistic duplex architecture, shown previously to exhibit ion channel activity^10^, we uncover changes in its structure *via* all-atom molecular dynamics (MD) simulations and experimental measurements. When introduced at the nicked position, cholesterol pulling towards the membrane can initiate unwinding of the double strand, which introduces flexibility that distorts the structure of the DNA forming the ion channel. This in turn affects the channel’s activity. Furthermore, we show that the distorting effect is hindered when hydrophobic anchors are introduced internally, linking the phosphate backbone, rather than at the terminal ends. The membrane activity of the non-nicked design - its pore stability and insertion efficiency - is significantly improved by this change in the construct design.

## Results and discussion

### Repulsive DNA-lipid interactions distort the molecular structure of a nicked duplex

We probe the influence of base pairing stability on membrane activity of synthetic DNA-based ion channels with three designs (Fig. 1a): DNA duplexes carrying two cholesterol moieties, differing in the distance between the modifications and the number of nicks. Two structures were comprised of four strands each, with cholesterols positioned at the nicks. Modifications were introduced symmetrically towards the center of the construct, separated by either 24 base pairs (bp) (≈ 8 nm) or 12 bp (≈ 4 nm). The third structure consisted of two strands, with cholesterols introduced as internal modifications, separated by 8 nm. We will refer to these three structures using the following notations: (distance between cholesterols) - (number of nicks), *i.e., 4nm-2x, 8nm-2x, 8nm-0x*. Figure 1a schematically shows the designs, alongside the chemical structure of the cholesterol linkage (Fig. 1b).

**Figure 1.**
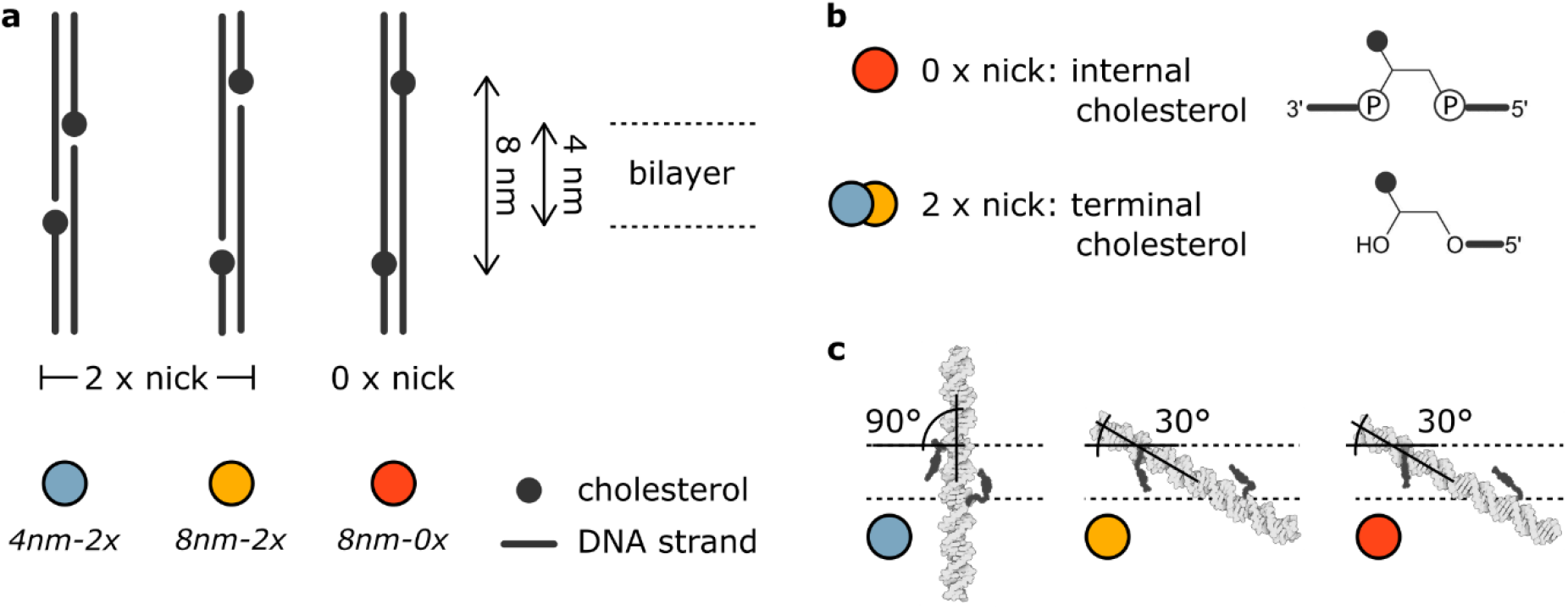
Schematic representation of the DNA constructs. (a) Sketch illustrating differences between the structures: distance between two cholesterol modifications and number of nicks in the modified positions. (b) The chemical linkage of introduced modifications: cholesterol in between two phosphates of a backbone in the absence of nicks or cholesterol at the terminal end, in the position of a nick. (c) Snapshots from the MD simulations, illustrating the initial configuration of duplexes in the lipid bilayer (represented as dashed lines).

Details of DNA sequences can be found in Supplementary Fig. 1 and Supplementary Table 1. NUPACK^35^ analysis of the sequences shows a lower (60-70%) base pair formation probability for two nucleotides adjacent to each nick, as compared with a 100% probability for base pairs mid-strand (Supplementary Fig. 2). The three structures were assembled using commercially available strands (SI, Section S1) and characterized with respect to their folding yield and stability (UV-vis absorbance profile and gel electrophoresis, Supplementary Discussion 1), as well as their bilayer interactions (imaging their attachment to model bilayers *via* fluorescent confocal microscopy, Supplementary Discussion 2). Their membrane insertion was investigated through transmembrane current measurements, reported in detail in the next section. Additionally, all-atom MD simulations provided a microscopic perspective on the structure of the DNA-lipid assemblies.

Based on the three designs, we have built three all-atom models, each containing a DNA structure embedded in a DPhPC lipid bilayer and surrounded by electrolyte solution. Considering that the thickness of the membrane is around 4 nm^36–38^, both *8nm* structures were inserted in a tilted conformation (under a 30° angle to the bilayer) in order to place both cholesterol anchors within the volume occupied by the lipid membrane. These structures mimic the orientations of natural membrane-associated helices, which span a wide range of tilt angles^39–41^. Driven by the hydrophobic mismatch, strongly resembling the principles guiding the orientation of the DNA helices introduced here, peptides can reach tilt angles even substantially larger than 30° ^39,40,42^. The *4nm-2x* structure, with its cholesterol spacing adjusted to the bilayer thickness, was oriented perpendicularly to the bilayer. Starting from these initial configurations (Fig. 1c), we performed 1 μs long equilibrium MD simulation for each system, as described in the methods section (SI, Section S2). Figure 2a shows the conformation of the duplexes towards the end of the simulation. Supplementary Movies 1-3 illustrate the dynamics of *4nm-2x, 8nm-2x* and *8nm-0x* DNA duplexes in the membrane.

**Figure 2.**
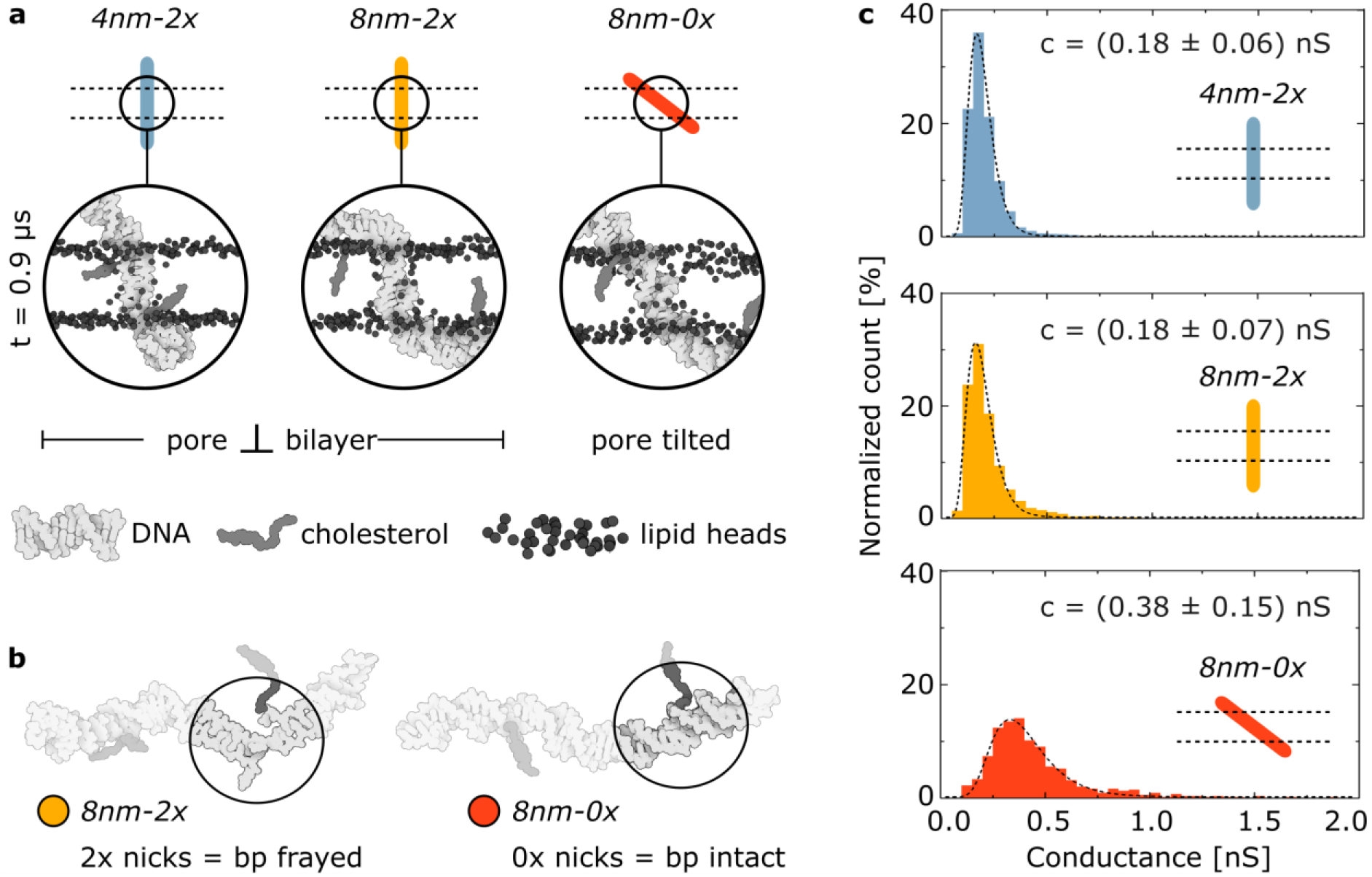
The effects of repulsive DNA-lipid interactions on the molecular structure of the duplexes. *(a) Sketches and respective frames from all-atom MD simulations at t = 0.9 μs. Lipid tails, ions and water molecules are not shown for clarity. (b) Snapshots from the MD simulations highlighting frayed base pairs at the position of one of the nicks in 8nm-2x structure, while the respective base pairs of 8nm-0x stay intact. (c) Histograms of reported signals from experimental ionic current measurements for all three structures. The dashed lines represent lognormal fits, with the conductance peak values stated on each plot. The error values represent standard deviation (SI, Section S2). The data was collected from three independent experiments, each 2 hours-long (total of 6 hours for each structure)*. N*_4nm-2x_* = 4287, N*_8nm-2x_* = 4857, N*_8nm-0x_* = 542.

To accommodate the *8nm* spacing between the cholesterols, a large part of the hydrophilic DNA was initially exposed to the hydrophobic core of the membrane. Due to this unfavorable orientation, the tilt of the *8nm-2x* design was observed to change over the course of the MD simulation. The fraying of the base pairs at the nicks allowed for straightening of the tilted structure, while keeping both hydrophobic molecules anchored in the bilayer’s core (Fig. 2a). Importantly, the *8nm-0x* design was initially oriented in the membrane in a similarly tilted manner, and presumably subject to the same deforming forces. Yet, its final conformation was much less distorted than its nicked analogue, with intact base pairings. We attribute this to the stability of base pairing in the absence of the nicks: non-nicked structure does not fray, and therefore it does not have the additional flexibility that enables bending of the nicked duplex. Figure 2b highlights the differences between base pairing in the analogous positions in the *8nm-2x* and *8nm-0x* constructs, as observe in our all-atom simulations.

The starting configuration of the *4nm-2x* design already has the optimal orientation of the DNA duplex with respect to the membrane to minimize the duplex’s exposure to the hydrophobic core of the bilayer. Even though this orientation does not change over the course of the MD run, the part of the helix embedded in the lipid membrane is observed to unwind (Supplementary Fig. 3a). Similar increase in the rise is reported for the *8nm-2x* structure. The natural distance between adjacent base pairs (3.4 Å)^43^ within the membrane increases to approximately 4.5 Å for both nicked constructs. As the unwinding minimizes the number of nucleotides confined within the hydrophobic core of the bilayer, we deduce that DNA-lipid repulsion is stronger than the forces introducing the helical twist to the double-stranded DNA (dsDNA). Note, that while changes in the rise are also reported for the non-nicked duplex, the latter does not find a stable, unwinded conformation within the timescale of the simulations, unlike the two nicked constructs (Supplementary Fig. 3b). Comparison of the duplex structures presented in Fig. 2b hints at a possible relationship between frayed base pairs at the nick and the distortion of the helical twist. This correlation has been observed in nature, where the helicase enzyme was shown to initiate dsDNA unwinding at the nick’s position^44,45^.

Summing up, structural distortions can be expected for nicked DNA nanostructures interacting with a lipid membrane. Such distortions can directly impact the membrane activity, which in our simulations was strongly correlated with the size of the pore in a lipid bilayer^13,24^ induced by the DNA. The pore surrounding the tilted *8nm-0x* nanostructure facilitated a larger transmembrane flux of water molecule (Supplementary Fig. 4) and a larger ionic current (obtained using SEM method^46^, Supplementary Fig. 5), than the pores formed by the nicked *4nm-2x* and *8nm-2x* constructs.

Experimental results allowed further confirmation of the notions observed *via* simulations. The insertion of the structures was studied by measuring the ionic current through DPhPC membranes in a 0.5 M KCl solution, while applying voltage of 50 mV (details of the transmembrane current measurements in the SI, Section S3). Applied voltage aids insertion of negatively-charged DNA, however we have previously showed that the insertion of these structures also occurs spontaneously^10^. In the absence of membrane-spanning structures, no current was recorded, while upon DNA insertion an increase in the signal was readily identified. Automatic analysis of the current traces by the Axon™ pCLAMP™ software suite (Clampfit) enabled finding all single-channel signals. Assuming ohmic behavior, each signal was attributed to the formation of a pore of defined conductance. Figure 2c shows histograms of the collected signals for each structure.

The conductance values obtained for the two constructs with nicks are indistinguishable, suggesting similar pore size (lognormal fit peaks at 0.18 ± 0.06 nS and 0.18 ± 0.07 nS for *4nm-2x* and *8nm-2x*, respectively). The comparable values obtained for nicked structures correspond with the simulation results, showing similar orientation of the formed pores (Fig. 2a). However, despite comparable magnitudes, the dwell times of the experimentally detected insertion events differ considerably (distribution presented in Supplementary Fig. 6). For *4nm-2x* we observe narrowly distributed, short signals, while durations of *8nm-2x* steps have a much wider range. Even though the two nicked duplexes form pores of similar sizes, their behavior in the membrane differs, and is determined by the distance between cholesterol modifications.

Importantly, the non-nicked *8nm-0x* structure has much wider conductance distribution, peaking at roughly twice the value: 0.38 ± 0.15 nS, suggesting a bigger pore size. Following the results of the simulations, we attribute the difference between the two *8nm* constructs to the mode of cholesterol linkage: non-nicked duplex induces a tilted pore, because, unlike its nicked analogue, it does not gain the flexibility (by fraying) to straighten the structure up. By removing the nicks, the architecture of the duplex in the membrane remains as designed, without significant distortion.

The conductance values found here are higher than the ones reported previously for similar structures, yet still of the same order of magnitude (reported 0.1 nS for a solution of twice higher conductivity^24^). The difference may result from chemical variations between constructs, since the previously reported duplex was anchored in a membrane using six porphyrin modifications (logP in a range [8.9, 11.8]^47^, as compared with logP = 7.11 for cholesterol (ChemAxon, chemicalize.com)), resulting in a strongly hydrophobic environment in the pore, which in turn causes more disrupted ion flow, as we showed before^10^.

On the other hand, the conductance measured *via* simulations for *4nm-2x* and *8nm-0x* (~ 0.17 nS and ~ 0.40 nS, respectively) agree with values obtained from experiments (~ 0.18 nS and ~ 0.38 nS, respectively). The similar conductance values reported from experiments and simulations suggest that indeed *8nm-0x* forms bigger pores, rather than inserts as a dimer, which could also explain its stronger signal. Conductance recorded in simulations for *8nm-2x* (~ 0.31 nS), even though smaller than for a non-nicked analogue, does not reach values similar to the *4nm* structure as in the experiments (~ 0.18 nS). However, it was to be expected, as the difference between timescales of simulations and experiments will play a more significant role in the case of the *8nm-2x* construct. The latter design was initially incorporated in the bilayer in a tilted conformation, and it requires time for the change in conformation. This change happens in nanoseconds and even though it may not be relevant in millisecond scale of experiments, it consists a large part of the simulation time. The final orientations of the remaining two structures do not differ as substantially from their initial ones, thus the simulated conductances reflect well the long timescale behavior detected experimentally for both *4nm-2x* and *8nm-0x*. In all cases the simulations qualitatively agree with experimental observations, and provide an insight into the likely molecular phenomenon responsible for our results. Through these outcomes, we confirmed our initial assumptions of the importance of base pairing stability and cholesterol linkage on the membrane activity of DNA-based transmembrane nanostructures.

### Improving insertion and pore stability by reducing the structure’s degrees of freedom

Collected current traces for the three duplexes show a transient ion flow, as presented in Fig. 3a (additional examples in Supplementary Fig. 7), which in turn suggests transient membrane-spanning: we determine each step as the structure going in and out of the bilayer. The duplex constantly changes its orientation and shape, to find the energetic minimum in the interplay between the cholesterol pulling towards the membrane and the DNA pulling away from it. By removing the nicks from the design, we effectively reduce its degrees of freedom, thus the structure has less ability to adapt. The control structure - non-inserting, single-cholesterol duplex - produces no signal, as expected (example of full-duration traces in Supplementary Fig. 7).

**Figure 3.**
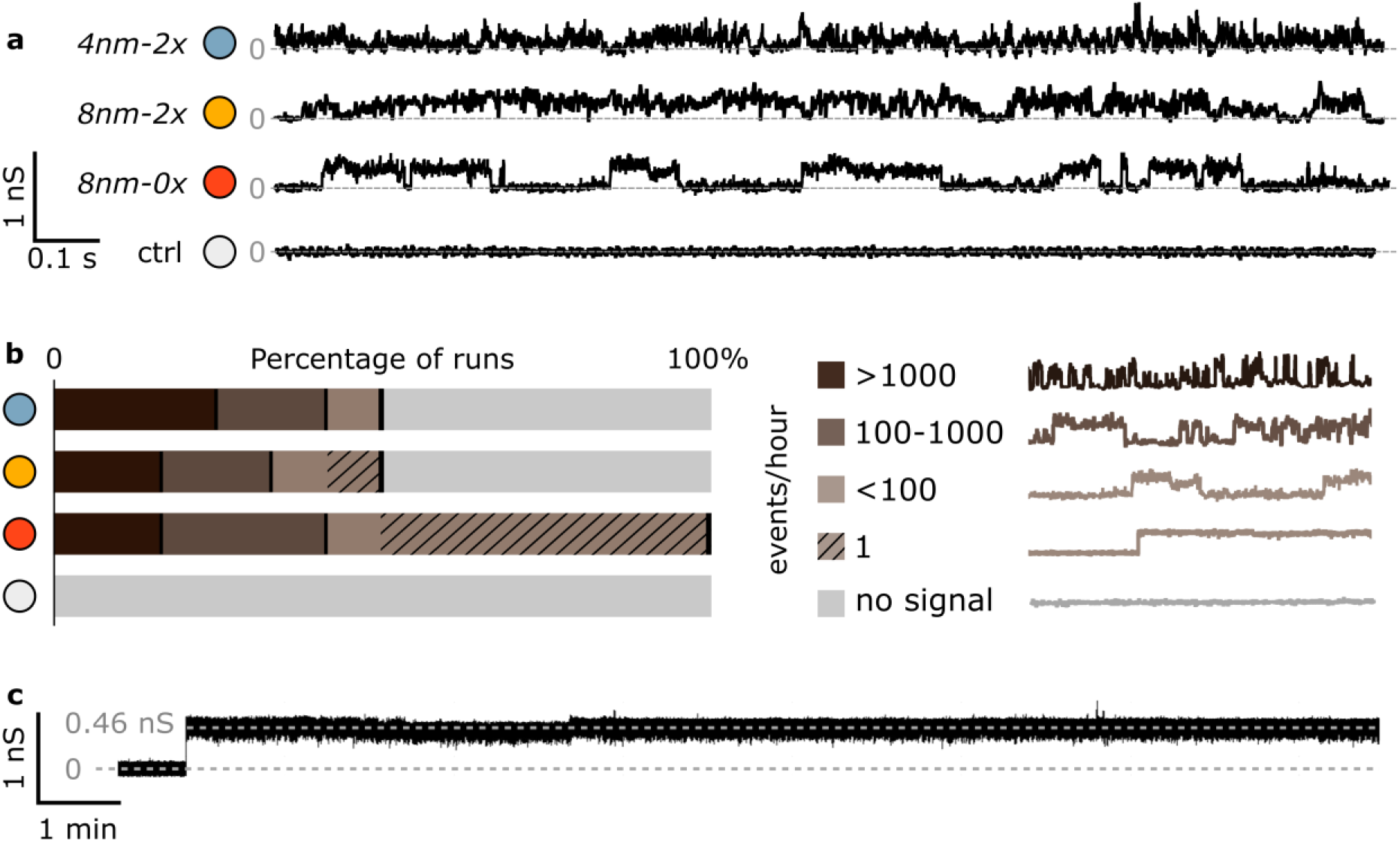
Improving pore stability and insertion efficiency by removing nicks from the design. (a) Examples of current traces collected for studied constructs. The “control” structure (ctrl, white) is a non-inserting duplex with a single cholesterol modification. (b) Plot illustrating insertion efficiency of studied structures, reporting percentage of runs with any signal above the noise threshold (± 0.05 nS). Twelve runs per structure, each at least 0.5 h-long. (c) An ionic trace showing a long-lasting signal, reported for the 8nm-0x structure. Further examples in Supplementary Fig. D3.1.

Consistent with this notion, for the *8nm-0x*, we observe more well-defined steps in the measured current. In contrast, the traces recorded for analogous nicked structure consist of much more varied signal, and the determination of the start and end points of the event is not trivial. This observation show that it is easier to clearly define two states - inserted and not inserted - for the *8nm-0x* structure containing no nicks. Meanwhile, the structure of nicked *8nm-2x* constantly changes by fraying bases and flexing at the nick’s position. A continuous alteration of the pore results in spikes and crooked steps in the current traces, for both nicked duplexes.

Despite its favorable cholesterol spacing, recordings for *4nm-2x* structure do not indicate a stable pore formation. In fact, the collected dwell times distribution of registered signals (Supplementary Fig. 6) shows that membrane-spanning events last short even compared with both *8nm* duplexes. This suggests that the distance between duplex’s end point and the cholesterol modification plays an important role in the insertion process.

The traces presented in Fig. 3a are examples of experiments with a detected DNA-induced signal attributed to pore formation (the non-inserting control returns the background current only). However, signals indicating ion channel activity were not always observed, even in the presence of inserting duplexes. We conducted twelve runs for each structure, each at least 0.5 h long., For illustration, the same number of runs with a signal above the noise level (> 0.05 nS) for every structure is plotted in Fig. 3b. As expected, no signal was induced by the non-inserting control structure. While the insertion of nicked constructs was only reported in 50% of the experiments, the presence of the non-nicked duplex *8nm-0x* led to transmembrane ionic currents in all bilayer experiments. We hypothesize that the stable positioning of insertion-driving cholesterol anchors contributes to the membrane-spanning process. When adjacent base pairs are not subject to fraying, the hydrophobic modifications are more likely to facilitate the formation of a DNA-induced pore.

Additionally, the majority of *8nm-0x* runs featured single or few steps of an exceptionally long duration, on a time scale of minutes. Such signals were observed almost solely for the non-nicked *8nm-0x* constructs, again hinting at the correlation between the stability of base pairing and pore formation. One representative current trace of a long-lasting signal is shown in Fig. 3c. More examples of long steps can be found in the Supplementary Discussion 3, alongside further analysis of recorded current traces and examples of high conductance steps and multiple insertions. Thus, the results presented in Fig. 3 indicate that the lack of nicks can have a strong stabilizing effect on the activity of membrane-spanning constructs. Decreasing the construct’s degrees of freedom resulted in a higher insertion efficiency, as well as better pore stability of the non-nicked structure - both highly desirable features of transmembrane protein mimics.

## Conclusions

We have investigated the effects of nicks in DNA-built transmembrane structures on their membrane interactions. Mindful of base pairs’ instability at the terminal ends of the strands, we studied the behavior of constructs modified with cholesterol either at the position of the nicks or introduced internally, with no discontinuities present in the double strand. The repulsive DNA-lipid interactions, combined with cholesterol’s membrane affinity, are strong enough to distort the molecular structure of the nicked construct by causing base pair fraying at the backbone discontinuities. The fraying introduces flexibility in the structure not accounted for by the design. However, in the absence of nicks base pairs adjacent to cholesterol are more stable and the DNA structure at equilibrium remains as designed.

Our analysis revealed other noticeable differences between nicked and non-nicked structures. We observed higher stability of the DNA-induced pores in the absence of nicks. Additionally, the insertion efficiency of these constructs was significantly improved. We attribute these effects to the correlation between the stability of hydrophobic anchors’ positioning and the membrane activity of nucleic acid nanostructures. With this we take steps towards understanding the process of membrane-spanning by hydrophobically-modified DNA, and we expect that further studies will elucidate a complete picture of the underlying molecular processes. Applying the insertion- and stability-improving modifications to existing larger designs can potentially change the perspectives of DNA-based protein mimics^13,31,32^, as well as nanopores for nucleic acid sequencing and analysis^48,49^.

By considering stabilization of base pairs, and the resulting limitations on the degrees of freedom of the DNA-built structures, we overcame distorting DNA-lipid interactions. Robust structural design is of particular significance for building biomimicking devices, where targeted environment features a vast range of complex chemical compositions. To that end, our DNA duplexes are inspired by natural transmembrane proteins that can tilt their transmembrane helices to undergo conformational transition affecting their activity, which can depend on the hydrophobic mismatch between the protein and the bilayer^50–52^. Further studies, comparing non-nicked DNA duplexes with various distances between cholesterols can shed more light on the importance of constructs’ orientation within the bilayer, as well as contribute a library of protein mimics with different tilt angles.

Even though the tilt of the membrane protein’s helix may vary, its helical *structure* stays unchanged in a wide range of hydrophobic mismatch arrangements^53^. In nature, the thickness of the bilayer is adjusted to proteins’ membrane-spanning domain^54^. In contrast, DNA duplexes, particularly the two nicked constructs, display significant changes to their helical structure upon membrane insertion, as we have shown through simulations (Supplementary Fig. 3). We suggest that the reported unwinding of the double strand has a potential to be utilized as a signaling pathway. For example, we speculate that a membrane-tethered DNA duplex could be designed to mimic a Talin mechanosensing protein^55^, whose functionality is based on transitions between stretched and unstretched conformation. While force sensing DNA nanodevices have been reported previously^56,57^, the unwinding observed here can also potentially be used to detect structural changes occurring within the bilayer. This notion is again inspired by nature, benefitting from the sensitivity of DNA’s helical structure for recognition purposes^58–60^.

Here, we showed that the idea of avoiding discontinuities in DNA strands can help preserve the designed shape, as well as improve pore-forming activity of the transmembrane structures. Applying our observations to a more complex design may result in a molecular machine that not only efficiently self-inserts into lipid membranes, but also performs in a stable and more controllable manner – an essential step towards building robust synthetic transmembrane structures. Arguably the most remarkable advantage of DNA nanoengineering is the ease of controlling molecular structure at nanoscale, thus, it is especially important to consider all relevant interactions including base pairing and fraying.

## Acknowledgements

D.M. acknowledges funding from the Winton Programme for the Physics of Sustainability and the Engineering and Physical Sciences Research Council (EPSRC, project ref. 1948702). S.E.S. acknowledges funding from the EPSRC Cambridge NanoDTC, EP/S022953/1. U.F.K. acknowledges funding from an ERC consolidator grant (DesignerPores 647144). A.A. and H.J. acknowledge support from the National Science Foundation USA (DMR-1827346). The supercomputer time was provided through the XSEDE allocation grant (MCA05S028) and the Leadership Resource Allocation MCB20012 on Frontera of the Texas Advanced Computing Center.

## Supplementary Data

Supplementary figures alongside the description of materials and methods used in this work are available free of charge in the attached document. Supplementary movies SM1, SM2 and SM3 show a 1 μs long all-atom MD simulation trajectories of the *4nm-2x, 8nm-2x, 8nm-0x* constructs, respectively. The complementary DNA strands are shown using turquoise and orange spheres and the backbone of DNA is shown in tubular representation. The lipid bilayer is shown using turquoise lines and a representative carbon atom (C22) of the lipid headgroup is shown as turquoise and yellow spheres to distinguish the upper and lower leaflets respectively. The cholesterol molecules are shown in green; water and ions are not shown for the sake of clarity.

---

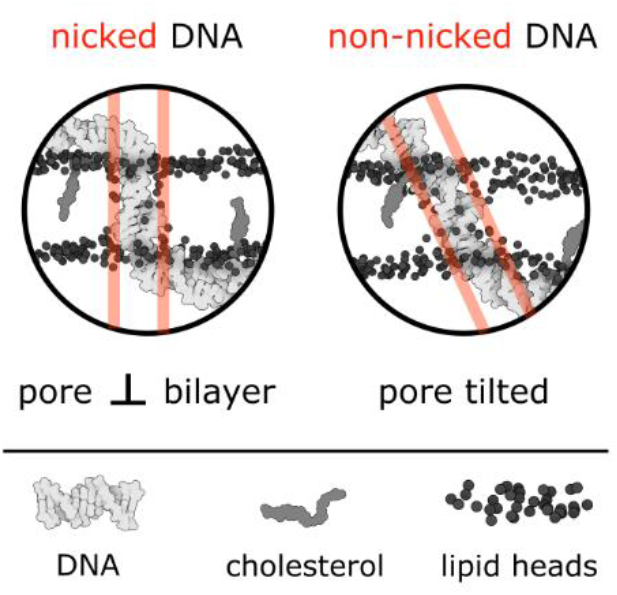
Table of Content graphic: Protein-mimicking DNA nanostructures can have their membrane activity improved by eliminating discontinuities (nicks) from the design. In the absence of nicks, cholesterol-driven bilayer insertion of DNA is more efficient, the formed pores more stable, and the molecular architecture of the construct does not get distorted. This allows control over helix’s tilt within the bilayer.

