## Supplementary figures, materials and methods for "Membrane activity of a DNA-based ion channel depends on the stability of its double-stranded structure"

[1] School of Engineering, École Polytechnique Fédérale de Lausanne, Route Cantonale, 1015 Lausanne, Switzerland

**S1 DNA nanostructure assembly**

All the reagents used in this work were acquired from Sigma Aldrich, unless stated otherwise. Each single strand was analysed using the NUPACK suite[1], in order to prevent formation of secondary structures, and to ensure sufficient yield of folding. For sequences see Supplementary Table 1. Oligonucleotides modified with an internal cholesterol were obtained from Eurogentec, while unmodified strands and end modifications (TEG (triethylene glycol)-cholesterol anchors, Cy3 labels) were provided by Integrated DNA Technologies, Inc. All the strands were dissolved to a final concentration of 100 µM: unmodified ones in IDTE buffer (10 mM Tris, 0.1 mM EDTA (Ethylenediaminetetraacetic acid), pH 8.0) and the modified in Milli-Q purified water. Strands were then stored at 4 °C, except for dye-modified ones, which were stored at -20 °C.

In order to fold the designed structures, the strands were mixed to a final concentration of 1 µM in TE4 buffer (10 mM Tris, 1 mM EDTA, 4 mM $Mg^{2+}$, pH 8.0), with cholesterol-modified strands heated beforehand at 70 °C for 10 min. DNA duplexes were heated up to 80 °C and then cooled down to 20 °C with 4 °C/min rate. Folded structures were all stored at 4 °C.

### S2 All-atom MD simulations

All MD simulations were performed using NAMD2[2]. The all-atom models of the 48 bp DNA duplexes having the same sequence used in experiments (Supplementary Table 1) were created using the NAB module of AMBERTOOLs[3]. A cholesterol molecule was covalently conjugated to the end of a strand using a triethylene glycol (TEG) linker, as described previously[4]. The force-field parameters for the cholesterol molecule with the linker were obtained from the CHARMM General Force Field (CGenFF) webserver[5]. The attachment points for the cholesterol molecules on the opposite strands of the duplex were separated either by 24 (*8nm-2x*) or 12 (*4 nm-2x*) bp, corresponding to approximately 8 and 4 nm, respectively (Supplementary Table 1). To obtain the no-nick variant of the *8 nm* design (*8nm-0x*), we created a custom patch and used it with the psfgen plugin of VMD[6] to make the DNA backbone continuous at the position of the cholesterol conjugation.

Each DNA construct was inserted into a pre-equilibrated patch of 1,2-diphytanoyl-sn-glycero-3-phosphatidylcholine (DPhPC) lipid bilayer membrane. In order to place both cholesterol anchors within the volume occupied by the lipid membrane, the *4nm-2x* design was inserted in a perpendicular conformation to the lipid bilayer whereas the *8nm-2x* and *8nm-0x* designs were inserted with a 30° tilt with respect to the bilayer (see Fig. 1c of the main text). All lipid molecules located within 3 Å of the DNA were removed. $Mg^{2+}$- hexahydrates were added near the backbone of the DNA to neutralize its negative charge, as described previously[7]. The resulting system was solvated with TIP3P water molecules[8] using the Solvate plugin of VMD[6]. Sodium and chloride ions were added to produce a 500 mM solution using the Autoionize plugin of VMD. A few additional $Mg^{2+}$– hexahydrates and chloride ions were added to result in the 4 mM bulk concentration of $MgCl_2$. The final systems measured approximately 13 x 23 x 13 $nm^3$ and contained approximately 346,000 atoms.

The assembled systems were subjected to energy minimization using the conjugate gradient method to remove the steric clashes between the solute and solvent. Following that, we equilibrated the lipid molecules around the DNA for 20 ns, while harmonically restraining all the non-hydrogen atoms of DNA using a spring constant of 1 kcal $mol^{-1}$ $Å^{-2}$. Subsequently, we removed the harmonic restraints and performed 50 ns equilibration while maintaining the hydrogen bonds between the complimentary base-pairs of DNA using the extrabond utility of NAMD. Finally, we removed all the restraints and performed 1 µs production simulation of each system in a constant number of atoms (N), pressure (P = 1 bar) and temperature (T = 298 K) ensemble.

All MD simulations were performed using periodic boundary conditions and the particle mesh Ewald (PME) method to calculate the long range electrostatic interactions[9]. The Nose-Hoover Langevin piston[10] and Langevin thermostat were used to maintain the constant pressure and temperature in the system. CHARMM36 force field parameters[11] described the bonded and non-bonded interactions between DNA, lipid bilayer, water and ions. An 8-10-12 Å cutoff scheme was used to calculate van der Waals and short range electrostatic forces. All simulations were performed using a 2 fs time step to integrate the equation of motion. SETTLE algorithm[12] was applied to keep water molecules rigid, whereas RATTLE algorithm[13] constrained all other covalent bonds involving hydrogen atoms. The coordinates of the system were saved at an interval of 20 ps. The analysis and post processing of the simulation trajectories were performed using VMD[6] and CPPTRAJ[3] whereas an online Fortran program Illustrator was used to visualize the structures[14].

**S3 Ionic current measurements**

Ionic current measurements were carried out using solvent-containing membranes. Hexadecane (1% in pentane) was used to coat both sides of a hole (Ø = 0.15 mm) in the foil dividing *cis* and *trans* chambers of the Teflon cuvette. After 5 minutes of incubation, 700 µL of 0.5 M KCl, 25 mM HEPES (4-(2-hydroxyethyl)-1-piperazineethanesulfonic acid), pH 7.0 was added to each chamber. 5 µl of 5 mg/ml DPhPC lipids (1,2-diphytanoyl-sn-glycero-3-phosphocholine, Avanti Polar Lipids) in pentane were added dropwise to each side, then the whole solution was gently pipetted up and down until the membrane was formed. Current data was acquired at a sampling rate of 5 kHz using Axopatch 200B amplifier. After membrane formation, DNA structures and o-POE (octyl-polyoxyethylene) surfactant were added to the *cis* side at the final concentration of 10 nM and 0.01 % respectively, and the ionic current under 50 mV voltage across the membrane was recorded. The experiments were repeated twelve times for each construct, and run for at least half an hour each. Clampex and Clampfit softwares were used to gather and analyse the data. "Single channel search" tool of Clampfit was used to automatically detect events reported in this work. Each dataset was analysed using the same settings: ignoring effects < 10 ms and only detecting single-level changes, with the level initialized at 10 pA (0.2 nS). Assuming an ohmic behaviour of the formed pores, conductance (c) was reported as recorded current (I) by voltage (V):

$$c = \frac{I}{V} \tag{1}$$

Lognormal distribution curves were fitted to the obtained histograms, following function (2).

$$y = y_0 + \frac{A}{\sqrt{2\pi}wx} e^{\frac{-\left[ln\frac{x}{x_c}\right]^2}{2w^2}} \tag{2}$$

where $y_0$ – offset, $x_c$ – center, $w$ – log standard deviation, $A$ – area.

The standard deviation was calculated using formula (3). Both formulae reported by Origin software, used for plotting the data.

$$\Delta y = e^{\ln(x_c)+0.5w^2}\sqrt{e^{w^2} - 1} \tag{3}$$

Origin software was also used to analyse dwell time data of the collected events. Function (4) was used to calculate kernel density ($k$).

$$k = \frac{1}{n}\sum_{i=1}^{n} \frac{1}{\sqrt{2\pi}w} e^{\frac{-(x-vX_i)^2}{2w^2}} \tag{4}$$

where $x$ – analysed dwell time, $vX$ – distributed samples used as kernel centres, $n$ – size of vector $vX$, $vX_i$– ith element of vector $vX$, $w$ – bandwidth used as kernel scale.

**S4 Assessment of DNA constructs' temperature stability using UV-Vis absorption spectroscopy**

The folding and stability of the DNA constructs was assessed using a UV-vis spectrophotometer (Cary 300 Bio, Agilent); thermal studies were performed in order to obtain melting curves of the unmodified structures. 100 µl of 1 µM DNA sample folded in TE4 buffer were heated from 10 to 90 °C, with a heating rate of 1 °C/min. Absorbance spectra were collected at 260 nm, and the melting temperature was obtained from the median of the two linear regions (upper and lower). The results are presented in Supplementary Fig. D1.2. The data and its analysis was processed using Origin software for all measurements taken.

**S5 Native polyacrylamide gel electrophoresis (PAGE)**

Polyacrylamide gel electrophoresis was used to confirm the proper folding of DNA designs. The gels were prepared at a concentration of 10% polyacrylamide, 0.5x TBE (Tris, borate, EDTA) and with 11 mM $MgCl_2$. Addition of 0.01 vol% ammonium persulfate (APS) (10%) and $6.7 \times 10^{-4}$% N,N,N',N' Tetramethylethylenediamine (TEMED) were used to initialise polymerisation, which proceeded for an hour. 2 µl of a DNA sample was mixed with 0.4 µl of 6x loading dye (15% Ficoll R400, 0.9% Orange G diluted in Mili-Q water), and then 2 µl of sample were loaded into the well. GeneRuler Low Range ladder (Thermo Fisher Scientific Inc.) was used as a reference. The gel was run in a Mini-PROTEAN R Tetra Cell (Bio-Rad), in 0.5x TBE with 11 mM $MgCl_2$ at 100 mV for 90 min. After this time the gel was immersed for 10 min in GelRed (Biotium), in order to stain the DNA. The imaging was performed on a GelDoc-It TM (UVP). The results are presented in Supplementary Fig. D1.1. FIJI was used to process gel images[16].

**S6 Confocal microscopy imaging**

Vesicles were prepared with electroformation, as reported previously[4]. POPC (1-palmitoyl-2-oleoyl-glycero-3-phosphocholine) and NBD-PC lipids (1-palmitoyl-2-{6-[(7-nitro-2-1,3-benzoxadiazol-4-yl)amino]hexanoyl}-sn-glycero-3-phosphocholine), both acquired from Avanti® Polar Lipids, were used in a ratio of 200:1, with the final concentration of 5 mg/ml in chloroform. 600 µl of 1 M sorbitol in 200 mM sucrose was used as a buffer. The osmolality of the buffer was around 1200 mOsm, with all the dilution buffers used in the experiments adjusted accordingly. Since for cell plasma the osmolality ranges between 275 - 325 mOsm[17], therefore we do not claim a biological osmolality. All the buffers were adjusted to pH 7.5 (using sodium hydroxide and hydrochloride solutions) - the value within the acidity range observed in natural systems[18].

Confocal microscopy images were acquired on an Olympus FluoView filter-based FV1200F-IX83 laser scanning microscope using a 60x oil immersion objective (UPLSAPO60XO/1.35). Cy3 excitation was performed using a 1.5 mW 543 nm HeNe laser at 1% laser power, with emission collected between 560 and 590 nm. FIJI was used to analyse the images. Representative micrographs showing membrane attachment of 8 nm constructs is shown in Supplementary Fig D2.1.

FRAP measurements were performed with the field of view focused on the top of a GUV. Using the FRAP function of the microscope's software (tornado mode), a spot of Ø = 4 µm was bleached and the fluorescence recovery observed. 10 images were collected pre-bleaching. Bleaching was performed over 0.5 s with 99% laser power and the fluorescence recovery was recorded for 50 frames.
Collected recovery curves were fitted using exponential function (5).

$$I_t = A\left(1 - \exp\left(-\frac{t}{\tau}\right)\right) + I \tag{5}$$

where $I_t$– fluorescence intensity in time t, $A$ – fitting parameter, $I$ – final intensity after recovery, $\tau$ – recovery time constant.

$\tau$ was then used to calculate recovery half-time as in (6).

$$t_{\frac{1}{2}} = \tau ln2 \tag{6}$$

Which in turn enabled obtaining diffusion coefficient $D$ following the formula (7).

$$D = \frac{0.88r^2}{4t_{\frac{1}{2}}} \tag{7}$$

*r* – radius of bleached area.[19]

Box plots with the collected diffusion coefficient values, alongside representative fluorescence recovery traces, is shown in Supplementary Fig. D2.2.

**S7 Supplementary Discussions**

**Supplementary Discussion 1. Structures' folding and stability**

Polyacrylamide gel electrophoresis (PAGE) was performed to ensure proper folding of three duplexes (Supplementary Figure D1.1), while UV-vis spectrophotometry – collected absorbance at 260 nm in a 10-90 °C temperature range – was used to assess temperature stability, alongside folding yield of the constructs.

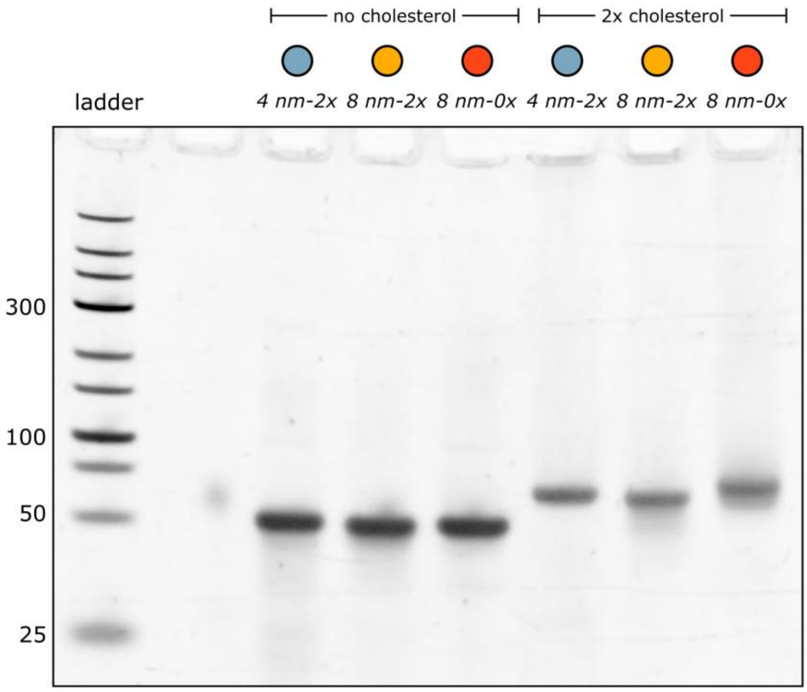

***Supplementary Figure D1.1.*** *Polyacrylamide gel electrophoresis analysis confirms proper folding of unmodified and cholesterol-modified duplexes. The lower intensity of bands featuring structures with hydrophobic anchors is attributed to their clustering, reducing their electromobility and causing them to remain in the well of a gel, and produce smeared bands.*

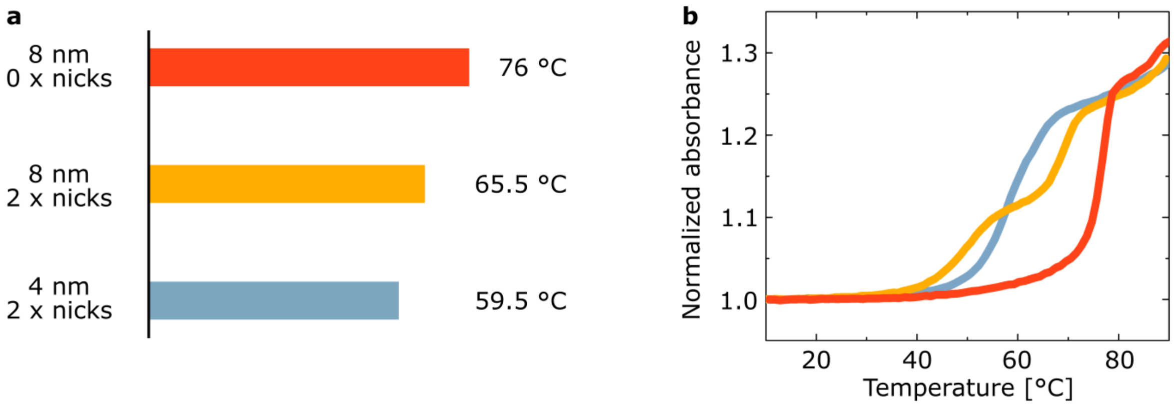

***Supplementary Figure D1.2***. *Results from melting unmodified DNA-constructs. (a) Melting temperatures were calculated from curves (b) obtained through spectroscopic measurements – absorbance at 260 nm collected in the presented temperature range.*

**Supplementary Discussion 2. Optical assessment of 8 nm structures' membrane interactions**

In order to confirm that cholesterol-modified duplexes are interacting with membranes, optical imaging through confocal fluorescence microscope was performed. Of particular interest were *8nm* structures and their comparison.

Duplexes modified with Cy3 fluorophore were incubated in the presence of POPC vesicles, and representative micrographs are presented in Supplementary Fig. D2.1a. Constructs were found to be coating the vesicles, indicating strong affinity towards lipid bilayers. Initially puzzling difference in the fluorescence intensity was proved to result from differences in the intrinsic fluorescence of each Cy3-modified strand, rather than differences in the membrane attachment efficiency, as showed by fluorimetric measurements presented in Supplementary Fig. D2.1b.

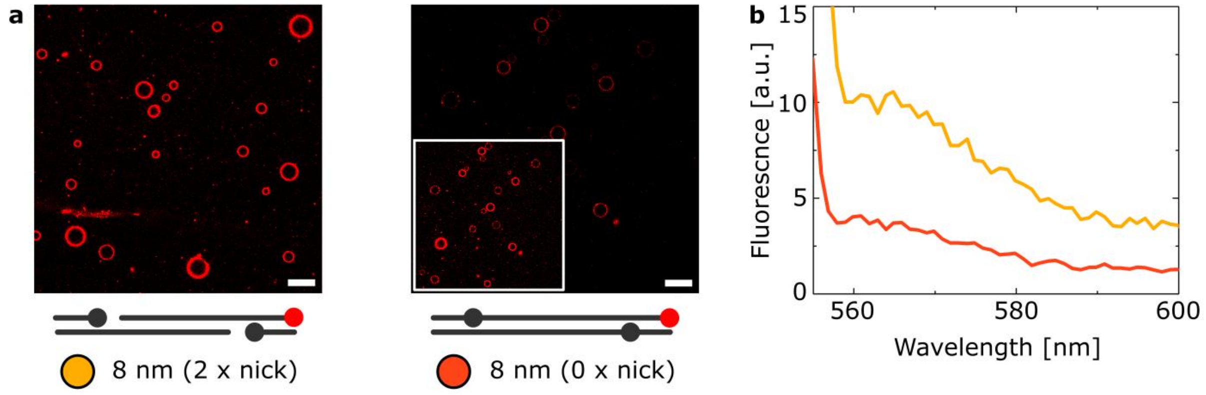

***Supplementary Figure D2.1.*** *Liposome coating by the 8nm structures. (a) Representative micrographs, showing Cy3-labelled DNA attachment to POPC vesicles. Scale bar: 20 µm. The inset on the graph corresponding to 8nm-0x shows the same area imaged with increased laser intensity. The lower fluorescence of 8nm-0x strands (with double modifications), responsible for the difference in recorded coating intensity, is confirmed by (b) fluorimetric measurements of Cy3-labeld strands of 8nm-2x (yellow, no cholesterol) and 8nm-0x (red, with cholesterol).*

FRAP measurements were performed on the DNA-coated vesicles, following Cy3 optical signal. Box plots of the calculated diffusion coefficient, alongside representative fluorescence recovery trace for *8nm-0x* and *8nm-2x* are presented in Supplementary Fig. D2.2. The results indicate that non-nicked construct diffuses in the membrane faster than its nicked analogue. Since the diffusion rate of cholesterol-tethered DNA has been reported to decrease with the increasing number of cholesterol anchors[20–23], the observed difference may result from the stronger anchoring of the nicked duplex. If the population of *8nm-2x* has both cholesterols stably positioned in a membrane, the anchors exert no force to overcome DNA-lipid repulsion, which hints on the reason behind its lower insertion efficiency.

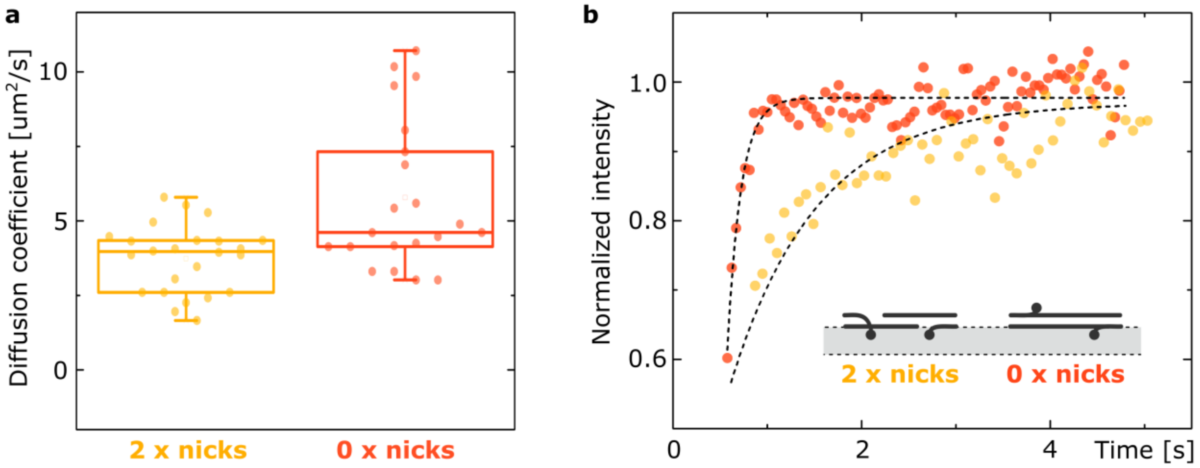

***Supplementary Figure D2.2.*** *Fluorescence recovery after photobleaching (FRAP) measurements results. Comparison of diffusion coefficient obtained for 8nm-2x and 8nm-0x structures in a form of a box plot (a), alongside representative fluorescence recovery traces (b). The inset in (b) presents hypothetical arrangement of the duplexes on the surface of the bilayer (grey), explaining the differences in their diffusion rate.*

**Supplementary Discussion 3. Further examples of observed events**

Apart from analysis of single-level changes in the current, multiple insertions were also observed for all three structures. With the exception of the *8nm-0x* construct, which often caused single, long-lasting steps (examples of which can be seen in Supplementary Fig. D3.1), upon a certain time (tens of minutes), multiple constructs were invariably seen to affect membrane's conductance. Representative examples of collected traces, alongside their all-point histograms are presented in Supplementary Fig. D3.2.

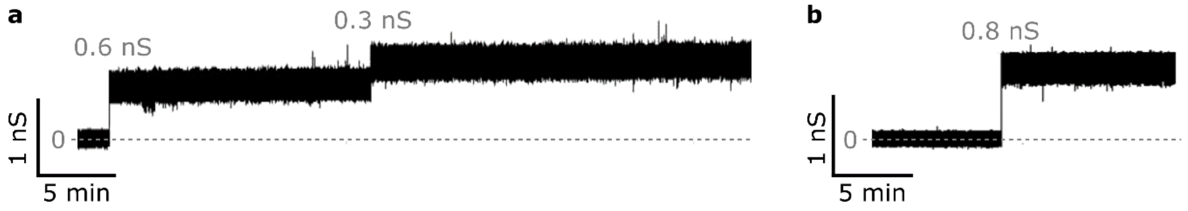

***Supplementary Figure D3.1*** *Further examples of long events observed in transmembrane current measurements.*

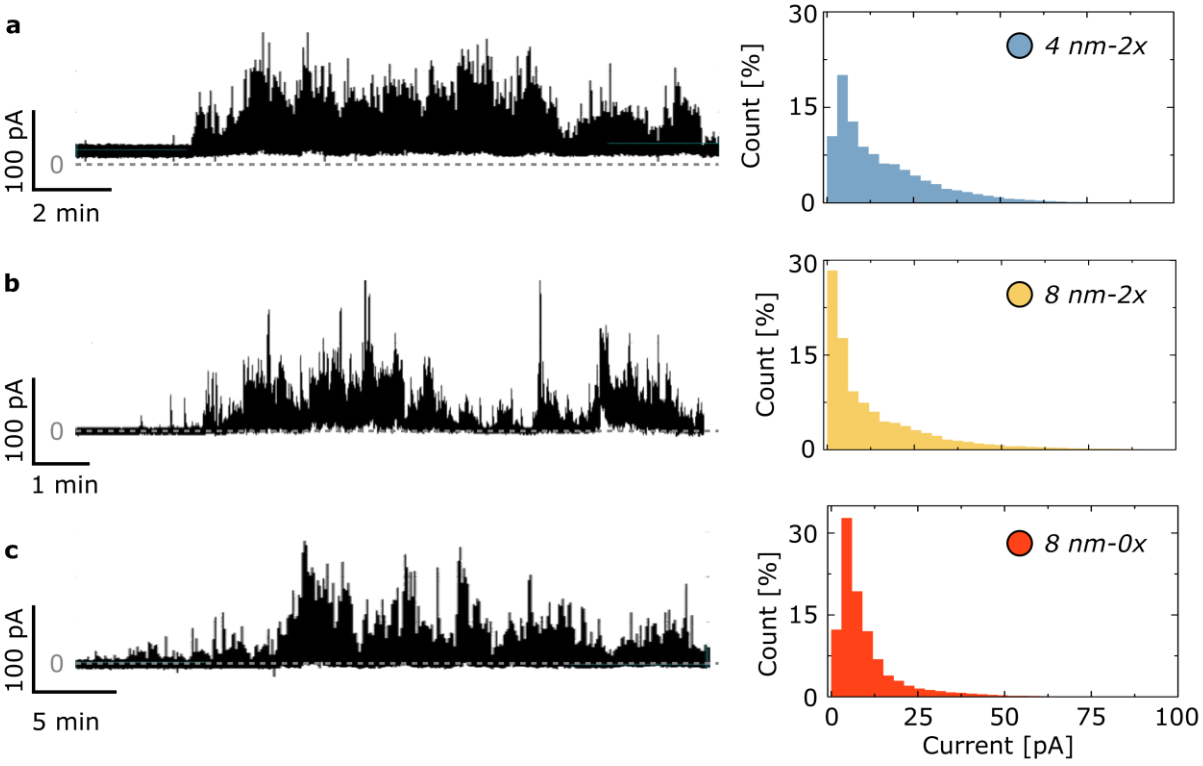

***Supplementary Figure D3.2.*** *Traces illustrating the long-term behaviour of studied constructs. The exemplary traces showing multiple insertions, with respective all-point histograms are shown for (a) 4nm-2x, (b) 8nm-2x, (c) 8nm-0x.*

However, due to the short dwell times, clear, discreet steps of multiple levels are rarely observed – even for the non-nicked structure. The data presented in the main text was usually extracted from the initial stages of the experiments, when well-defined events could be distinguished. Still, for non-nicked *8nm-0x* structure we did observe examples of multiple insertions, like the ones shown in Supplementary Fig. D3.3.

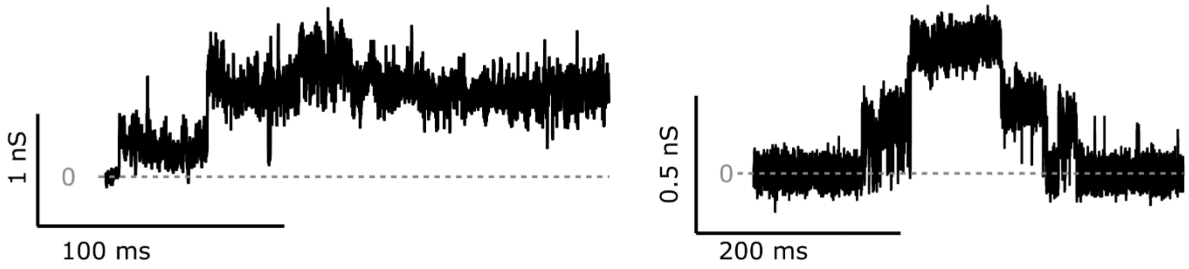

***Supplementary Figure D3.3.*** *Exemplary traces showing clear multiple insertions of 8nm-0x structure.*

Similarly, examples of high conductance steps (Supplementary Fig. D3.3.) are reported for *8nm-0x*, which we suggest may result from structure's clustering and inserting as a multimeric modular channel, that could be responsible for higher detected signals.

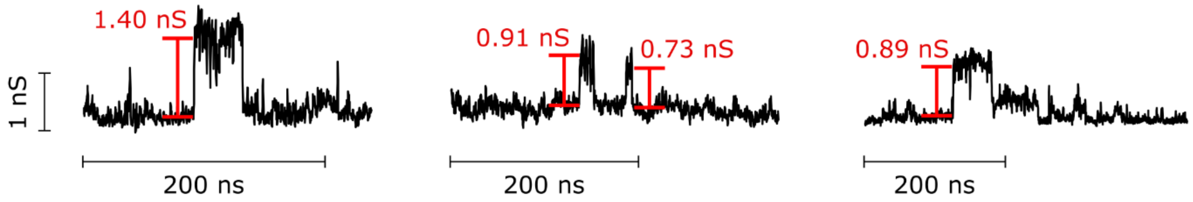

***Supplementary Figure D3.4.*** *Example traces showing high-conductance steps recorded for 8nm-0x construct.*

**S8 Supplementary Figures**

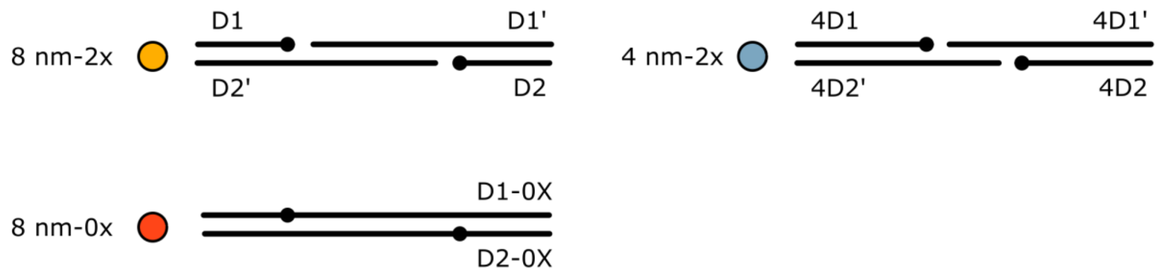

***Supplementary Figure 1*** *Schematic representation of the three designs used in this work. The sequences of labelled strands can be found in Supplementary Table 1.*

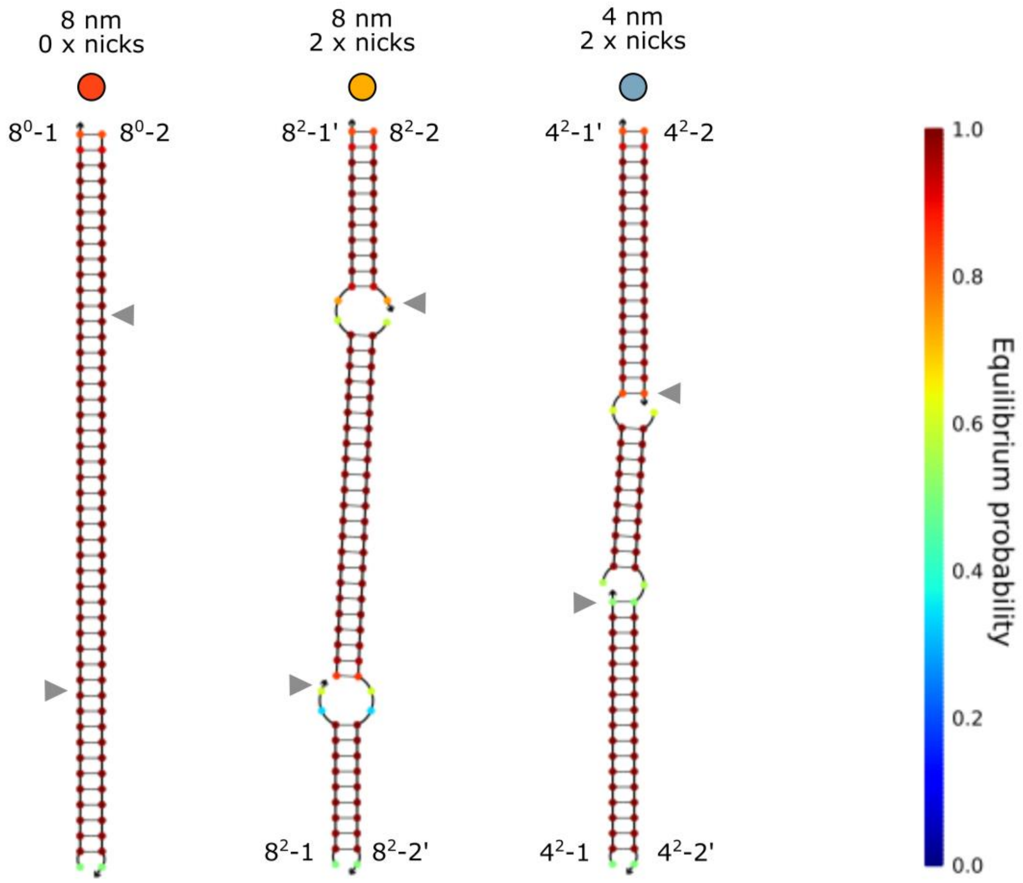

***Supplementary Figure 2.*** *NUPACK analysis of three designs used in this work. The colour map illustrates probability of forming base pairs. Grey arrows point at the positions of cholesterol in each construct.*

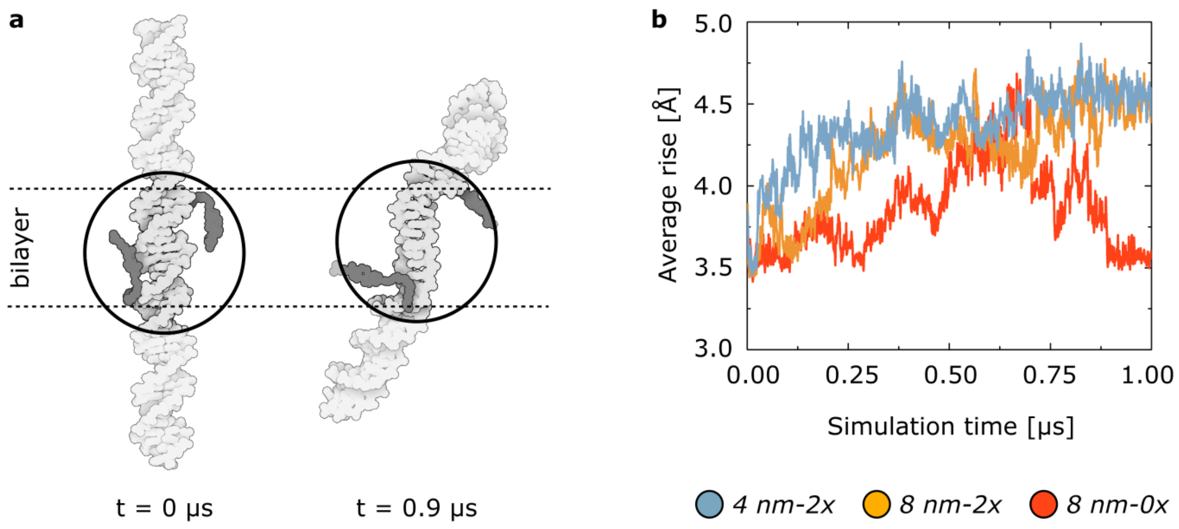

***Supplementary Figure 3.*** *Change to the DNA twist in the membrane-spanning domains, observed via simulations. (a) Snapshots from simulations of 4nm-2x in the bilayer, highlighting prominent change with respect to the double-helix conformation in the membrane-spanning region. (b) The average rise of the membrane-spanning base pairs of DNA (24 and 12 base-pairs for 8nm and 4nm constructs respectively) measured as a function of simulation time for the three structures. The 8nm-0x construct shows the less deviation from the ideal B-DNA structure as compared to the 8nm-2x and 4nm-2x constructs.*

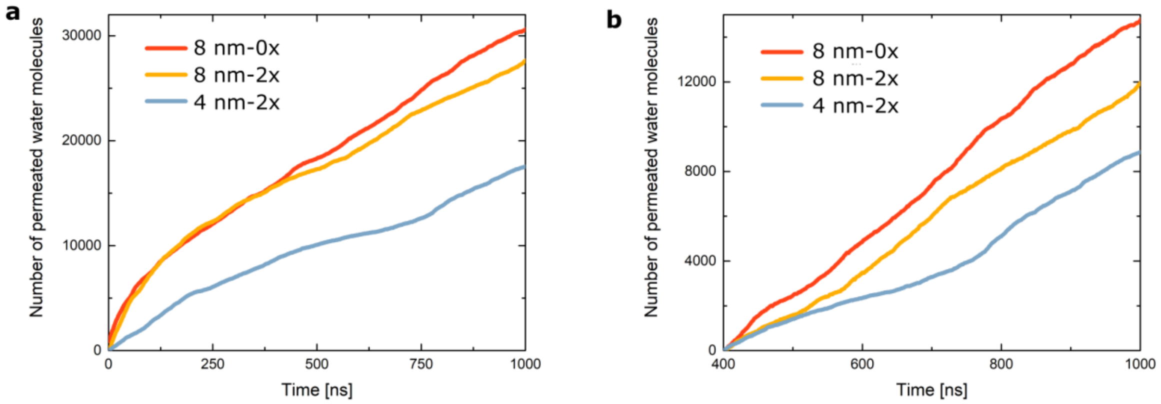

***Supplementary Figure 4.*** *Number of permeated water molecules observed in MD simulations: throughout the whole simulation (a), and normalized change after the first 400 ns (b).*

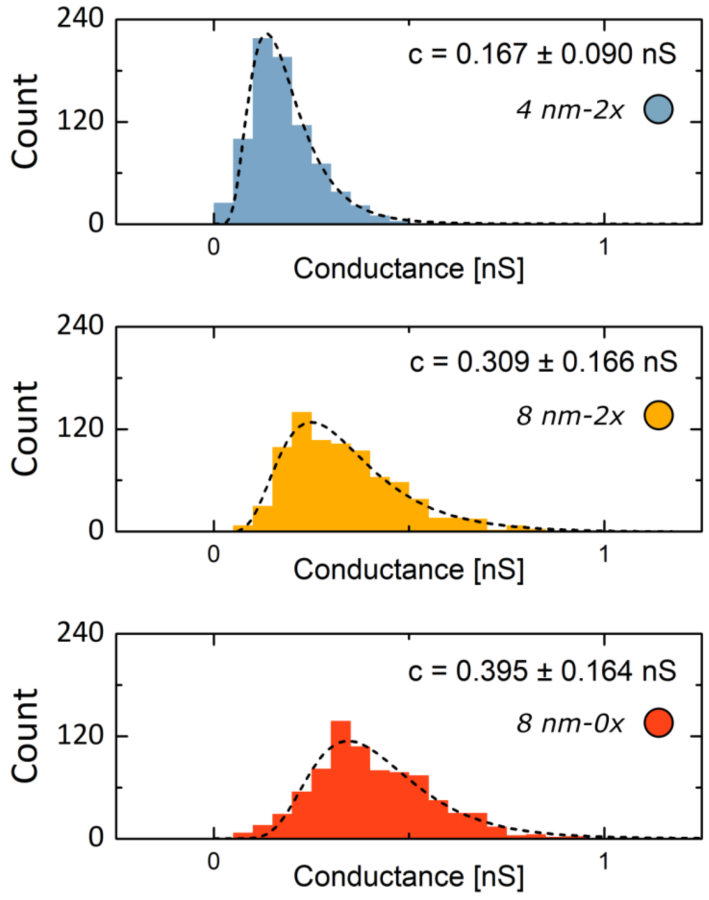

***Supplementary Figure 5.*** *All-point histograms of ion conductance recorded from the last 800 ns of the simulations. Dashed lines represent lognormal fit. Peak values, alongside standard deviation, are stated on each plot. The ionic currents were computed using the SEM method[15]. The 8nm-0x construct is found to conduct the ionic current the most, followed by 8nm-2x and 4nm-2x.*

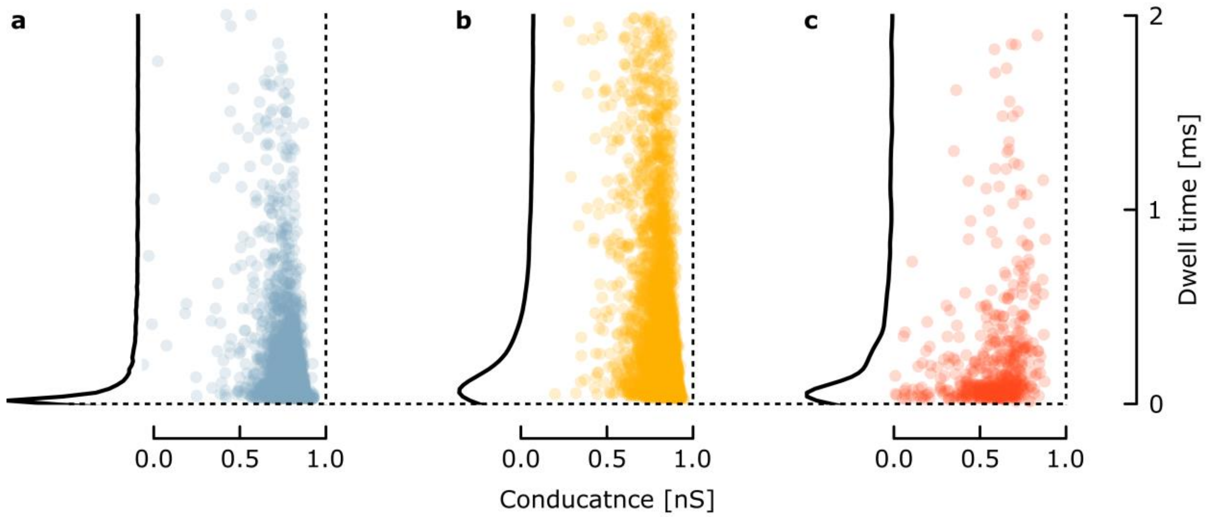

***Supplementary Figure 6.*** *Scatter plots of conductance vs dwell time of detected events (reported in histograms in Fig. 2c of the main text) for (a) 4nm-2x, (b) 8nm-2x, (c) 8nm-0x. The kernel distribution plot for each sample's dwell times is presented next to the respective scatter plot – the distribution clearly indicates the differences between the two nicked structures (4nm-2x and 8nm-2x), despite their conductances distributed similarly and their number of detected events comparable:* $N_{4nm\text{-}2x} = 4287$, $N_{8nm\text{-}2x} = 4857$, $N_{8nm\text{-}0x} = 542$.

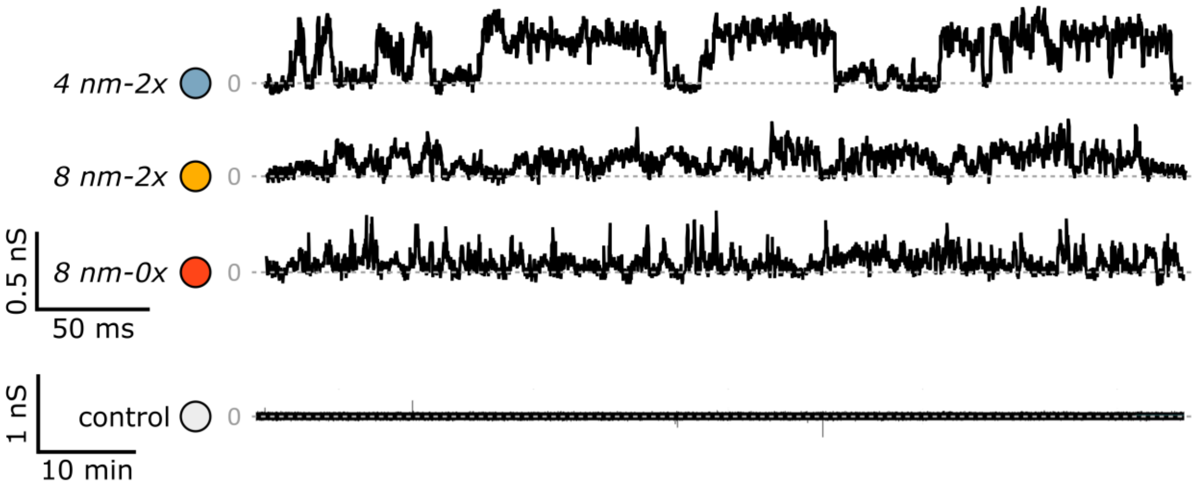

***Supplementary Figure 7.*** *Further examples of traces recorded via transmembrane current measurements. No signal was reported in control experiments: the single spikes in the current were among the short (< 10 ms) events, and therefore attributed to experimental artefacts (e.g. sensitive setup picking up movement in the laboratory).*

### S9 Supplementary Tables

***Table 1*** *DNA sequences used in this work. • represents the position of cholesterol. The schematic representation of the strands and the structures they form can be found in Supplementary Fig. 1.*

| *Structure* | *Strand* | *Sequence* | *Length* | *Modification* |
|---|---|---|---|---|
| 4 nm<br>2 x nick | $4^2$-1 | AGTAGTATCCATCATCGT• | 18 | 3’ cholesterol |
| | $4^2$-1’ | AGCTTTTTAAGTCATACATAGATTAGAGAG | 30 | 5’ Cy3 |
| | $4^2$-2 | CTCTCTAATCTATGTATG• | 18 | 3’ cholesterol |
| | $4^2$-2’ | ACTTAAAAAGCTACGATGATGGATACTACT | 30 | |
| 8 nm<br>2 x nick | $8^2$-1 | AGTAGTATCCAT• | 12 | 3’ cholesterol |
| | $8^2$-1’ | CATCGTAGCTTTTTAAGTCATACATAGATTAGAGAG | 36 | 5’ Cy3 |
| | $8^2$-2 | CTCTCTAATCTA• | 12 | 3’ cholesterol |
| | $8^2$-2’ | TGTATGACTTAAAAAGCTACGATGATGGATACTACT | 36 | |
| 8 nm<br>0 x nick | $8^0$-1 | AGTAGTATCCAT•CATCGTAGCTTTTTAAGTCATACATAGATTAGAGAG | 48 | int. cholesterol<br>5’ Cy3 |
| | $8^0$-2 | CTCTCTAATCTA•TGTATGACTTAAAAAGCTACGATGATGGATACTACT | 48 | int. cholesterol |
